# ViReMa: A Virus Recombination Mapper of Next-Generation Sequencing data characterizes diverse recombinant viral nucleic acids

**DOI:** 10.1101/2022.03.12.484090

**Authors:** Stephanea Sotcheff, Yiyang Zhou, Yan Sun, John E. Johnson, Bruce E. Torbett, Andrew L Routh

## Abstract

Genetic recombination is a tremendous source of intra-host diversity in viruses and is critical for their ability to rapidly adapt to new environments or fitness challenges. While viruses are routinely characterized using high-throughput sequencing techniques, characterizing the genetic products of recombination in next-generation sequencing data remains a challenge. Viral recombination events can be highly diverse and variable in nature, including simple duplications and deletions, or more complex events such as copy/snap-back recombination, inter-virus or inter-segment recombination and insertions of host nucleic acids. Due to the variable mechanisms driving virus recombination and the different selection pressures acting on the progeny, recombination junctions rarely adhere to simple canonical sites or sequences. Furthermore, numerous different events may be present simultaneously in a viral population, yielding a complex mutational landscape. We have previously developed an algorithm called *ViReMa* (Virus Recombination Mapper) that bootstraps the *bowtie* short-read aligner to capture and annotate a wide-range of recombinant species found within virus populations. Here, we have updated *ViReMa* to provide an ‘error-density’ function designed to accurately detect recombination events in the longer reads now routinely generated by the Illumina platforms and provide output reports for multiple types of recombinant species using standardized formats. We demonstrate the utility and flexibility of *ViReMa* in different settings to report deletion events in simulated data from Flock House virus, copy-back RNA species in Sendai viruses, short duplication events in HIV, and virus to host recombination in an archaeal DNA virus.

## Introduction

Recombination is essential for virus evolution and adaptation. Homologous recombination allows for the reshuffling of single nucleotide variants (SNVs) among viral genomes or for the formation of chimeric viral genomes when they co-infect a single cell (1). Such homologous recombination events have been attributed to the emergence of outbreak strains of viruses including rhinoviruses and coronaviruses (2–4). Recombination events that give rise to new and culturable strains/species of virus can readily be identified though phylogenetic comparison of full-length consensus genomes. Such studies inform on historical recombination events that gave rise to novel and ‘successful’ emergent viruses that have survived selectivity filters.

Similarly, non-homologous recombination in viral genomes can give rise to diverse genetic species. These range from simple insertions and deletions, to more complex rearrangements of the virus genome as well as inter-genic recombination between different RNA virus species and/or their host. Deletion or insertion events can result in the formation of structural variants. For example, small 4-5 amino acid deletions are commonly seen surrounding the furin cleavage site of the SARS-CoV-2 spike protein after cell-culture passaging (5). Larger structural variants have been reported that remove or disrupt entire viral genes, such deletions in ORF 8 upon adaptation of SARS-CoV-1 to human transmission (6). Simple duplications of the viral genome are frequently observed during the intra-host diversification of HIV during anti-retroviral treatments. These duplications include short 9-21 AA stretches near the protease cleavage sites of GAG that result in altered processing kinetics of GAG by the HIV protease in the presence of protease inhibitors (7–11).

If the product of a non-homologous recombination event is not a viable viral replicon, then the product is termed a ‘defective’ viral genome (DVG) (12, 13) or ‘Defective RNA’. Generally, DVGs are unable to produce the viral proteins required for replication, participle assembly, or other aspects of the viral replication cycle. Nevertheless, DVGs can still be replicated and passaged in *trans* by the parental or ‘helper’ virus. DVGs sometimes have the ability to compete with or otherwise attenuate their parental ‘helper’ viruses though a variety of different proposed mechanisms such as sequestration of cellular viral cofactor, competition with their helper viruses for access to the viral polymerase or via strong immuno-stimulation. Such DVGs are termed as ‘*interfering*’. While the presence and interference of DVGs has long been well appreciated in cell-culture conditions (14), it is now emerging that the spontaneous emergence of DVGs during viral infections can have significant impacts on the outcomes of viral pathogenesis and vaccine design (15–17). The term ‘defective’ may therefore be somewhat misleading as it appears that such species may play an important role in the normal lifecycle of some viruses.

Non-homologous recombination may also give rise to intergenic recombination between two or more orthogonal templates. For example in Flock House virus, a bi-partite RNA virus from the *Nodaviridae* family, chimeric genetic species comprising fusion of the two genomic segments are spontaneously generated intracellularly during the replication cycle (18). Intergenic recombination can also occur between the viral genomes and their hosts. For example, fragments of human ribosomal protein mRNAs have been found to be integrated into the genomes of Hepatitis E virus (HEV) that were extracted from persistently infected patients. Interestingly, these virus-host chimeric strains displayed a growth advantage in cell culture (19, 20).

Next-Generation Sequencing (NGS) provides a high-throughput and quantitative tool to characterize the frequencies of different genetic products of viral recombination in complex virus populations. Canonical spliced or gapped sequence aligners such as *HISAT2* (21, 22) and *STAR* (23), are suitable to map simple deletion-type recombination events. There have also been reported tools that can discover and annotate different types of recombination events found within viral genomes including: Host-to-virus recombination/integration sites (*ViralFusionSeq* (24), *VERSE* (25)); copy-back and snap-back DVGs (*DI-Tector* (26), *VODKA* (27)); and intra-strain homologous recombination events (*Hexahedron* (28), *MosaicSolver* (29)). Many of these pipelines require prior knowledge/information to resolve recombination events and they make assumptions as to the ‘types’ of event that are found. However, due to the diverse and complex nature of viral recombination, the capture and quantification of the full range of genetic products of viral recombination requires a similarly diverse and versatile computational tool.

We previously reported an algorithm for detecting and counting recombination events found in next-generation sequencing analyses of viral genomes that we called ‘*ViReMa*’, (**Vi**ral **Re**combination **Ma**pper) (30). *ViReMa* bootstraps the *bowtie* or *bwa* short-read aligners (31, 32) to map short Illumina reads to a reference genome. For reads in which only a partial mapping is found, *ViReMa* extracts the unmapped portion of the read and re-maps this segment in a new mapping iteration. If the subsequent segment maps discontinuously, then *ViReMa* reports a recombination event. Importantly, the mapping of the subsequent segment is completely independent of the previous segment, which allows *ViReMa* to identify any kind of recombination event among either strand of any reference sequences provided (virus or host or otherwise). As a result, *ViReMa* can ‘agnostically’ report any complex recombination event.

*ViReMa* has been independently validated by Alnaji et al (33) and Boussier et al (34) who provided a series of condition and parameter optimizations using simulated and experimental data to map RNA recombination events in DVGs of influenza A and B. *ViReMa* has been used and validated to study viral recombination in range a of plant viruses (Polygonum ringspot virus (PolRSV) (35), Tomato yellow leaf curl virus (TYLCV) (36), Potato X Virus (37)), insect viruses (Flock House virus (FHV) (38), Cricket Paralysis virus (CrPV) (39)), arboviruses virus (Zika virus (ZIKV) (40), Chikungunya virus (CHIKV) (41, 42)), human pathogens (Influenza virus (43), Reovirus (44), MERS-CoV and SARS-CoV-2 (45, 46)), and retroviruses (Human Immunodeficiency Virus (HIV) (7)).

NGS technologies have advanced since the original inception of *ViReMa*, when NGS read lengths were seldom longer than 100nts. As a result, the original pipeline needed only tolerate a small, fixed number of mismatches within a mapped read segment. Here, we introduce a newer error-handling process that assigns recombination junction breakpoints once a threshold number of reference mismatches are encountered within a specific moving window, rather than simply counting the number of mismatches found in total across a mapped read segment. This is important due to the fact that NGS sequencing reads are now considerably longer than when *ViReMa* was originally conceived, for example, with MiSeq reads being up to 300bp in length. Furthermore, this allows *ViReMa* to accurately map reads from virus populations that have extensive intrahost diversity and so contain multiple minority variants relative to the reference genome.

We have also developed *ViReMa* to provide outputs in standardized bioinformatic conventions. Foremost, *ViReMa* provides alignments in SAM format (47) to allow visualization and integration with common downstream bioinformatic tools. Deletions and duplication events as well as more complex recombination events, such as copy-back or inter-genic fusions are reported in standardized BED or BEDPE formats (48). We have also added a simple GUI functionality, improving accessibility. To illustrate the broad functionality of *ViReMa*, here we analyze multiple different model virus systems including: Flock House virus (+ssRNA) for the detection of deletion-type recombination events; Sendai virus (-ssRNA) for the detection of copy-back DVGs, HIV (a retrovirus) for the detection of short InDels and genomic duplications, and STIV (dsDNA archaeal virus) for the detection of virus-to-host recombination junctions. Overall, *ViReMa* provides a validated and robust ‘one-stop’ solution to simultaneously capture the full-range of recombination events found within virus populations.

## Methods

### Description of Algorithm

*ViReMa* is a python3 script that boots-straps the small read mappers: *bowtie* (49) or *bwa* (50). Virus references are provided by the user in FASTA format (an index is automatically generated if one is absent), and the host reference must be a pre-built *bowtie* or *bwa* index. For short reference sequences without poly(A)-tails, additional A’s should be added to the end of the reference genome to allow *bowtie* to map reads that would otherwise overhang the end of the virus reference genome. In the absence of this, such reads would be rejected by *bowtie* preventing *ViReMa* from finding recombination events or chimeric reads that involve the very 3’ end of the viral genome. *ViReMa* uses *bowtie* (49) or *bwa* (50) to generate alignments of a fixed seed length starting from the left-most position of a sequence read. After this initial mapping, ViReMa tracks along the rest of the read segment following the mapped seed until a disqualifying mismatch is found, as would occur at the junction of a recombination event. Mismatches are not uncommon in short reads either due to errors in base-calling or biological variation. Therefore, 0-2 mismatches are tolerated in the mapped read before a junction is inferred. Single mismatches are not allowed to occur within ‘X’ nucleotides of the junction (the ‘X’ value can be specified in the command line, default value is 5).

With improvements seen in next-generation sequencing (NGS) and the common usage of longer Illumina reads up to 300nts in length, permitting only 2 mismatches per read became insufficient. Indeed, defining the number of allowed mismatches as a function of read-length lacks biological rationale as read-length is a purely technological choice and the number of mismatches is often determined by the base-calling error rate inherent to sequencing platforms and the intrahost diversity of the sample in question. Therefore, we have introduced a new mapping procedure. The basis of mapping is the same, whereby an initial seed is mapped using *bowtie* (49) or *bwa* (50), and then trailing nucleotides are assessed for evidence of a break. However, rather than counting the total number of nucleotide mismatches to locate a break-point, a break-point is elicited when a given number of mismatches are found within a defined window. These figures are given in the command line by an ‘Error-Density’ parameter. The default is (1,25); which means that up to 1 mismatch is tolerated within any 25 given nucleotide window. 2 mismatches within 26 nucleotide would be tolerated. 2 within 25 would invoke a recombination break-point at the most downstream mismatch as per the usual *ViReMa* protocol. This process has the primary advantage over the previous versions of allowing longer reads to be mapped before premature break-points are invoked due to simple mismatches.

### SAM output

We have substantially over-hauled and updated *ViReMa* in view of making the software more user-friendly and accessible. *ViReMa* standardizes the output, returning canonical SAM files with appropriate CIGAR 1.4 scores and associated SAM tags. The purpose of this was to allow users to visually inspect their alignments using common alignment visualization software such as the Tablet viewer (51) or the Integrative Genomics Viewer (IGV) (52, 53) and allow *ViReMa* to be implemented as part of routine RNAseq pipelines that require SAM files as an input.

For the most part, straight-forwardly mapped reads either with or without soft-pads simply use the output SAM information from the original *bowtie* or *bwa* alignment. Similarly, straight-forward recombination, deletion and/or splice events are reported as canonical alignments with either ‘D’ or ‘N’ nucleotides in between each mapped segment. The choice of whether to specify a gap as a deletion (‘D’) or a recombination/splice event (‘N’) is in the command line entry by the ‘--*MicroInDel*’ option, with a default value of 5 nts. Insertion events shorter than the value defined by the ‘*--MicroInDel*’ option are stored using ‘I’ in the CIGAR string. Longer insertions are stored as soft-pads (‘S’) in between two independently reported read segments. If longer insertions are in fact the result of duplication of a segment of the underlying genome reference (as illustrated in **Figure 2**), these can be reported as insertions in a single entry of the SAM file rather than two segments by invoking the ‘*--back-splicing*’ option. This allows these features to more easily be visualized in NGS alignment software.

*ViReMa* is also capable of mapping complex recombination events, such as when a number of inserted nucleotides that do not map to the reference genome are present between two or more mapped segments of a recombination event. Awareness of these types of complex recombination events is important in the context of understanding viral recombination, as mismatched nucleotides inserted into nascently replicated RNA/DNA strands may be a trigger for non-homologous recombination and template switching. These unmapped/unmappable nucleotides are stored as soft-pads (‘S’) in the output SAM file. These may occur at the end of mapped read segments or in between mapped red segments.

One of the advantages of using the *ViReMa* software is the ability to map unusual recombination events, such as trans-splicing events, inter-genic recombination, or large back-splicing or back-recombination events. Currently, there is no standardized system with which to report these types of alignments, without treating each mapped segments separately in the SAM file. In *ViReMa*, we do the same, treating these types of events as chimeric reads. Individual segments are specified using the “TC:i:n” and “FC:i:n” tags, and these reads are hard-padded at the segment break junctions.

### Annotation of Recombination events in Original Formats

All the original formats output of ViReMa have been retained to allow backwards compatibility, as previously described (30). Briefly, recombination junctions are output into text files with the simple format (for example): “*1102_to_1037_#_9*”. This means that there is a recombination junction with a donor site at nucleotide 1102 to an acceptor site at nucleotide 1037 in the reference genome and that 9 unique reads mapped over this junction. These junction annotations are listed underneath a header that describes the genomes that were mapped to either side of the junction (for example: “*FHV_to_FHV*”; or “*HIV_RevStrand_to_HIV_RevStrand*”).

However, we have made two simple additions to allow further scrutiny of these events. By invoking the “*-FuzzEntry*” option, the amount of microhomology found at the recombination junction is reported, for example: “*1102_fuzz-5_1037_#_9*”. This means that there is a recombination junction with a donor site at nucleotide 1102 to an acceptor site at nucleotide 1037 in the reference genome, that there are 5 nucleotides of microhomology at the recombination junction and that 9 unique reads mapped over this junction. We have also added a “*-ReadNamesEntry*” option that when invoked appends the read name of every unique read that mapped to each reported events after the annotation. This has applications when searching for specific reads in alignment visualizers to ‘spot-check’ and verify the authenticity of reported recombination events and may have applications in future single-cell sequencing pipelines that retain cellular barcodes and UMIs in the read names.

### Annotation of Recombination events in Standardized BED Formats

We have updated *ViReMa* to provide recombination event annotations using standardized BED and bedgraph formats to allow visualization and integration with only bioinformatic pipelines. Virus recombination events within viral genes are expressed using canonical BED format, similar to RNA splicing events or DNA recombination events, but with a minor alteration. As of *ViReMa* version 0.25, the first six columns are in BED6 format, with four additional bespoke columns:

1. Reference name taken from FASTA file of aligned genome;
2. Nucleotide Coordinate of donor site;
3. Nucleotide Coordinate of acceptor site;
4. Name of the type of event: “*Deletion*” if acceptor site is downstream of donor site; or “*Duplication*” if donor site is downstream of acceptor; “Ins:NNN” if a small insertion of ‘NNN’ nucleotides is found or if “*--BackSplice_Limit*” is set;
5. The number of mapped reads reporting this event;
6. The strand of the reference genome in which the event is found (“+” or “-”);
7. The total read coverage found at the donor site (includes all reads either with or without evidence of the recombination junction at this site);
8. The total read coverage found at the acceptor site (includes all reads either with or without evidence of the recombination junction at this site);
9. The nucleotide sequence found surrounding the recombination junction of the donor site;
10. The nucleotide sequence found surrounding the recombination junction of the acceptor site.

The inclusion of read coverage at the recombination junctions allows for an approximation of the recombination event abundance in the dataset. However, care must be taken when performing these calculations due to the unevenness and/or bias of read coverage at different sites in a viral genome which may result in different coverage at either recombination junction. For columns 9 and 10, the number of nucleotides reported upstream and downstream of the recombination junction is determined by the mapping seed used in the alignment (default is 25 nts) and the exact breakpoint mapped is indicated using a pipe: “|”. This provides a convenient output to assess for evidence of nucleotide bias at the sites of mapped recombination events as well as quickly assess whether there is micro-homology between recombination events, as is commonly seen, for example, in coronaviruses (45). If the ‘*--MicroInDel*’ option is specified in the command-line, an additional BED file is produced containing the annotations only for the microInDels.

For inter-genic and/or forward-to-reverse strand recombination events, we have adopted the BEDPE format (48). This format is illustrated in **Figures 1, 4** and **5** and can be used to describe end-to-end fusions of multi-partite RNA genes, inter-genic recombination events, copy/snap-back DVGs (examples shown in table) as well as virus-to-host fusion/chimeric events.

As of *ViReMa* version 0.25, the BEDPE format contains the following information:

1. Reference name of the first mapped reads segment
2. Coordinate of the nucleotide immediately upstream of the recombination/fusion junction of the first reference (given in column 1).
3. Coordinate of the nucleotide immediately downstream of the recombination/fusion junction of the first reference (given in column 1).
4. Reference name of the second mapped reads segment
5. Coordinate of the nucleotide immediately upstream of the recombination/fusion junction of the second reference (given in column 4).
6. Coordinate of the nucleotide immediately downstream of the recombination/fusion junction of the second reference (given in column 4).
7. Name of the type of event: “*Copy/Snap-Back*” if read segments map to different strands of same viral genome; “*Intergenic-Fusion*” if read segments map to different viral genes and or viral genomes; “*Virus-Host-Fusion*” if one read segments maps to a viral gene and one read segment maps to the host genome; and “*Gene-Fusion*” if read segments map to different host genes/chromosomes.
8. The number of mapped reads reporting this event;
9. The strand of the reference genome given in column 1 (“+” or “-”);
10. The strand of the reference genome given in column 4 (“+” or “-”).

**Figure 1.**
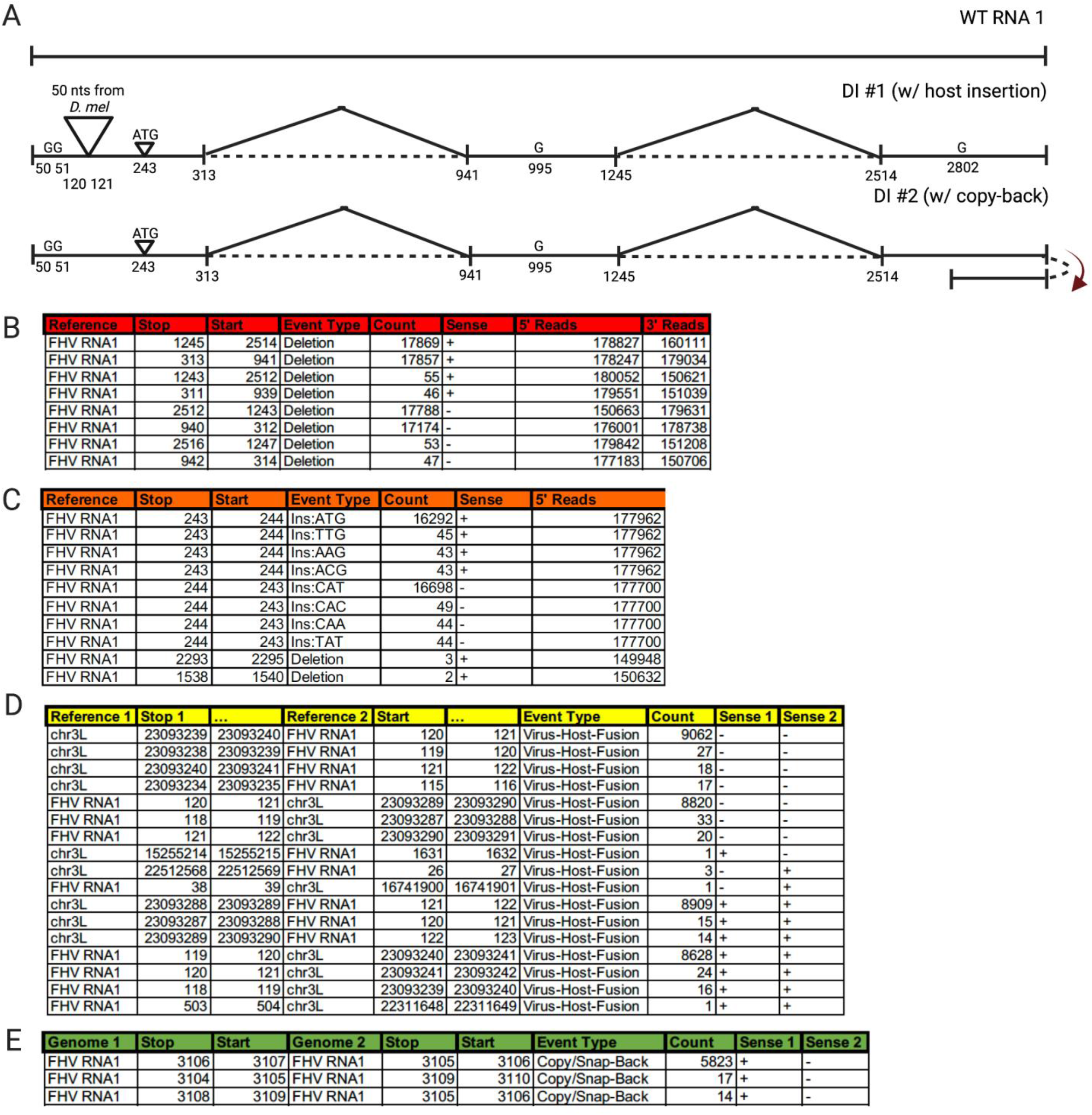
*ViReMa* analysis of simulated Flock House virus (FHV) data. **A)** Defective RNA1 (D-RNA1) genomes we constructed and simulated reads were generated using ‘ART’. Wild-type FHV RNA1 is depicted, with schematics of hypothetical D-RNAs shown below. D-RNA1 #1 includes a 50nt insertion from *D. melanogaster* as well as two deletions. D-RNA1 #2 includes the same deletion events and a duplication event at the 3’ end. **B)** *ViReMa* results of the simulated data are reported in the Virus_Recombination_Results.bed file. Output indicates the reference (in this case FHV has two references, one for each strand of its bi-partite genome), coordinates of the recombination break-points, the type of recombination event, the number of reads mapping to this even, the stranded-ness of the recombination event, and how many total reads at the 5’ and the 3’ end of the recombination junction. **C)** *ViReMa* results for simulated data found in the MicroRecombinations.bed file. These indicate the reference, the genome position the event occurred, the type of event, (Ins:XXX is an insertion of XXX nucleotides), coordinates of the recombination break-points, and number of reads at the 5’ and the 3’ sites of the recombination junction. **D)** *ViReMa* results for simulated data found in the Virus-to-Host-Recombination.BEDPE file. This displays the reference of the first genome mapped in reads and where that stops before mapping to the second reference at the “start” nucleotide position. The event type is also noted, as well as the count of that recombination event and the strandedness of genome of each mapped segment. **E)** *ViReMa* results for simulated data found in the Virus_Fusions.BEDPE file. Similar information is depicted as in the Virus-to-Host-Recombination output. Here we show the detection of the 3’ end duplication we included in our simulated dataset.

### Additional features and Optimizations

By profiling the rate/usage/time of each operation in the script, we determined that ~50% of the runtime can be attributed to each iterative *bowtie* alignment. As a result, datasets that have a large number of un-mappable reads or reads that would map to a reference that is not provided, will incur several rounds of futile iterative alignments. We have added a feature invoked using “--*MaxIters*” that allows users to limit the number of iterations that are attempted. This prevents long run-times associated with unmappable reads being repeatedly tested. This option may be important in scenarios where users do not have access to a complete assembled genome for the host and/or when the host material is the predominant species in the input RNA/DNA that was originally sequenced.

As unmapped reads are stored in temporary memory, there is a natural limit to the number of reads that can be aligned in one instance of *ViReMa*. As a result, very large datasets may get ‘stuck’ in the initial phases of the program’s initialization. We have added a feature invoked using “*--Chunk*” to allow users to analyze large datasets in ‘chunks’, whereby a user-defined number of reads are mapped in any given time. We set a default ‘chunk’ size of 1 million reads. This number was chosen as it typically corresponds to requirement for approximately 1-2Gb of RAM, which is easily accessible for most regular workstations, as which does not saturation the 2Gb limit imposed by *pypy*.

### Simulated data

The ART suite of tools for the simulation of NGS reads was used to generate datasets of read of synthetic recombination events and Defective-RNAs (D-RNAs) of Flock House virus (54). Parameters used were 37’000 X coverage over each of the two simulated D-RNAs and 300’000X over the wild-type virus. We generated individual FASTQ files for each synthetic FASTA reference file and reads were concatenated from multiple FASTQ files in ratios stipulated in the main text to generate single FASTQ files used in the *ViReMa* analysis. An error profile reflecting the HiSeq 2500 was imposed for reads 100 bases in length, yielding ~10M reads.

### Data Alignments and Visualization

All data were aligned to the virus genomes either with or without the host genome in parallel using *ViReMa* version 0.25 (30) using parameters and command-lines described in the main text for each virus sample. Output SAM files were transformed and sorted into BAM files using *samtools* (47) and coverage plots were obtained using *bedtools (48)*. Read alignments were visualized using *Tablet (51)*.

## Results

### Flock House virus recombination in simulated

We generated two artificial Defective-RNAs (D-RNAs) based upon recombination events previously seen in Flock House virus (FHV) as well as imaginary events designed to challenge and test our platform. These templates are illustrated in **Figure 1**. The first D-RNA contained two deletion events, a simple 3nt insertion, multiple point mutations as well as an insertion of a 50 nt fragment of host mRNA (chr3L:23’086’340-23’086’389 from *dm6*) at nt120 of the viral genome. The second D-RNA, instead of containing this host insertion event, contained a fusion-type recombination event (akin to a copy-back RNA) whereby the 3’ end of the viral genome is fused to the reverse sense of the 3’-most 41 nts of the genome. Using these input reference sequences, we generated simulated reads using ‘ART’ tools (55) requiring approximately 37’000X coverage over each simulated D-RNAs (74,000X total for both D-RNAs) and 300’000X over the wild-type virus. An error profile reflecting the HiSeq 2500 was imposed, yielding 10,935,310 reads.

We ran the *ViReMa* pipelines using standard parameters: (--X 3 --Defuzz 0 --Host_Seed 25 - BED --MicroInDel_Length 5) mapping to both a padded FHV genome and *Drosophila melanogaster* (*dm6)* genome. The resulting BED files produced are shown in **Figure 1B-C** and **SData 1**. As can be seen, each of the simulated recombination events are successfully detected with no erroneous mappings. The correct deletion recombination events at 313^941and 1245^2514 are found, as is a small ATG insertion between nts 243 and 244. The expected host insertion (virus-host fusion type) event was successfully detected, as reported using the BEDPE output **Figure 1D**. Both virus-to-host junctions were found corresponding to both ends of the insertion and with reads mapping in both the negative and positive sense orientation. Finally, the hypothetical positive to negative sense recombination event at the 3’ end of the viral genome (illustrated in **Figure 1A**) was also captured, as reported using the BEDPE output **Figure 1E.**

As the coverage of simulated reads over the defective genomes is approximately 37’000X for each D-RNA, we should therefore expect to find approximately 74’000 or 37’000 reads that map over either the shared or unique (respectively) synthetic recombination junctions illustrated in **Figure 1A**. However, recombination junctions cannot be found in the extremities of individual sequence reads as there are not enough bases available to unambiguously map either side of a recombination junction. With a seed length of 25nt and with 100nt long reads, there are only 49 potential ‘cutting sites’ (30) per read across which recombination junctions can be detected in a single read. Therefore, we can maximally only expect to find 36,260 or 18,130 reads that map over either the shared or unique (respectively) synthetic recombination junctions.

As seen in **Figure 1C**, *ViReMa* found 35,031 and 35,657 reads (combination of both positive and negative orientation) mapping to the deletion recombination events 313^941 and 1245^2514 respectively, yielding ~97% mapping efficiency. The imperfect sensitivity is due to the presence of single nucleotide mismatches that are found near recombination junctions. While these reads can be found in the output SAM file, *ViReMa* does not annotate these reads as containing recombination events if mismatches are found near the putative recombination junction at a distance defined by the optional --X parameter (set to 3 above). This value can be adjusted to increase sensitivity, but at the cost of introducing false positive events (43). As the host insertion event was only present in one of the synthetic D-RNAs, we would expect approximately 18,130 reads to map to either side of the insertion event. As seen in **Figure 1D**, we detected 17,729 and 17,690 (combination of both positive and negative orientation) mapping to each end of the host insertion event, again yielding ~97% mapping efficiency.

As the duplicated, negative-sense portion of this event was only 41 nts in length, this restricts the number of the 100nt longs reads that can map to this region as *bowtie* does not allow reads to overhand the ends of a reference genome. Therefore, with a seed length of 25 nts there are only 16 (41 – 25) possible cutting-sites remaining in any read across which this junction can be detected. From an expected coverage of 18’500X, we would therefore expect approximately 5,920 reads. ViReMa found the copy-back-like recombination event at the 3’ end of the viral genome with 5,823 reads **(Figure 1E)**, therefore yielding an efficiency of ~98%.

### Duplications in the GAG region of the HIV genome

In a previous study, we reported the analysis of covariation of amino acid and nucleotide variants within a large cohort of HIV-1 and antiretroviral treated (ART) patients who were part of the U.S. Military HIV Natural History Longitudinal Study (56, 57). We used *ViReMa* to detect insertions, deletion, or recombination events in HIV obtained from patient blood samples during ART. To begin, processed reads were mapped to the reference HIV-1 genome (**Figure 2A, B**) (NL4-3) using *ViReMa* (BED files are provided in and **SData 2)**. We invoked the *--back-splicing* option to report short insertions and duplications in the HIV genome, as described in *Methods*. This revealed a large number of recombination events occurring close to the protease cleavage sites between p17 (matrix) and p24 (capsid) proteins, as well as in the PTAP region of the p6 protein (**Figure 2A**). These events are largely characterized by in-frame duplications ranging from 3 to 24 nucleotides in length. The most common HIV sequence alterations resulting from resistance to inhibitor treatment in patients group into three HIV genomic regions and are (**Figure 2A**): 1) small duplications at the ‘AQQA’ motif found 13 amino acids upstream of the p17/24 protease cleavage site; 2) ‘QSRPE’ duplications two amino acids downstream of the p7 (nucleocapsid)/p6 (p1p6) protease cleavage site; and 3) ‘PTAP’ duplications 10 amino acids downstream of the p7/p6 protease cleavage site. Importantly, these duplications have been previously observed, and have been characterized as a response to anti-viral drug treatment (57–60). There is some variation in the exact nature of these duplications, varying in length from 3 to 24 nucleotides, sometimes containing inexact duplications that introduce variant amino acids at the duplication site. Nonetheless, the same or closely similar events were seen in multiple different patient samples, indicating that these duplications were a response to a common evolutionary selection pressure. In **Figure 2A** and **B,** we illustrate where AQQA and PTAP duplications are occurring in the context of the HIV genome and what these duplications are at the nucleotide level for two examples: a ‘PTAP’ duplication with coordinates 2157^2143 and an ‘AA’ duplication with coordinates 1148^1143. Note that in the third time point (Feb 2001) the 1148^1143 is replaced with 1150^1145, shown in the table in **Figure 2C.** The gray dotted line in panel **Figure 2D** illustrates when the patient was switched from indinavir (IDV) to another anti-retroviral – nelfinavir (NFV). After switching from IDV to NFV, the patient showed a substantial increase in the abundance of the ‘AA’ duplication and only a small increase in the PTAP duplication.

**Figure 2.**
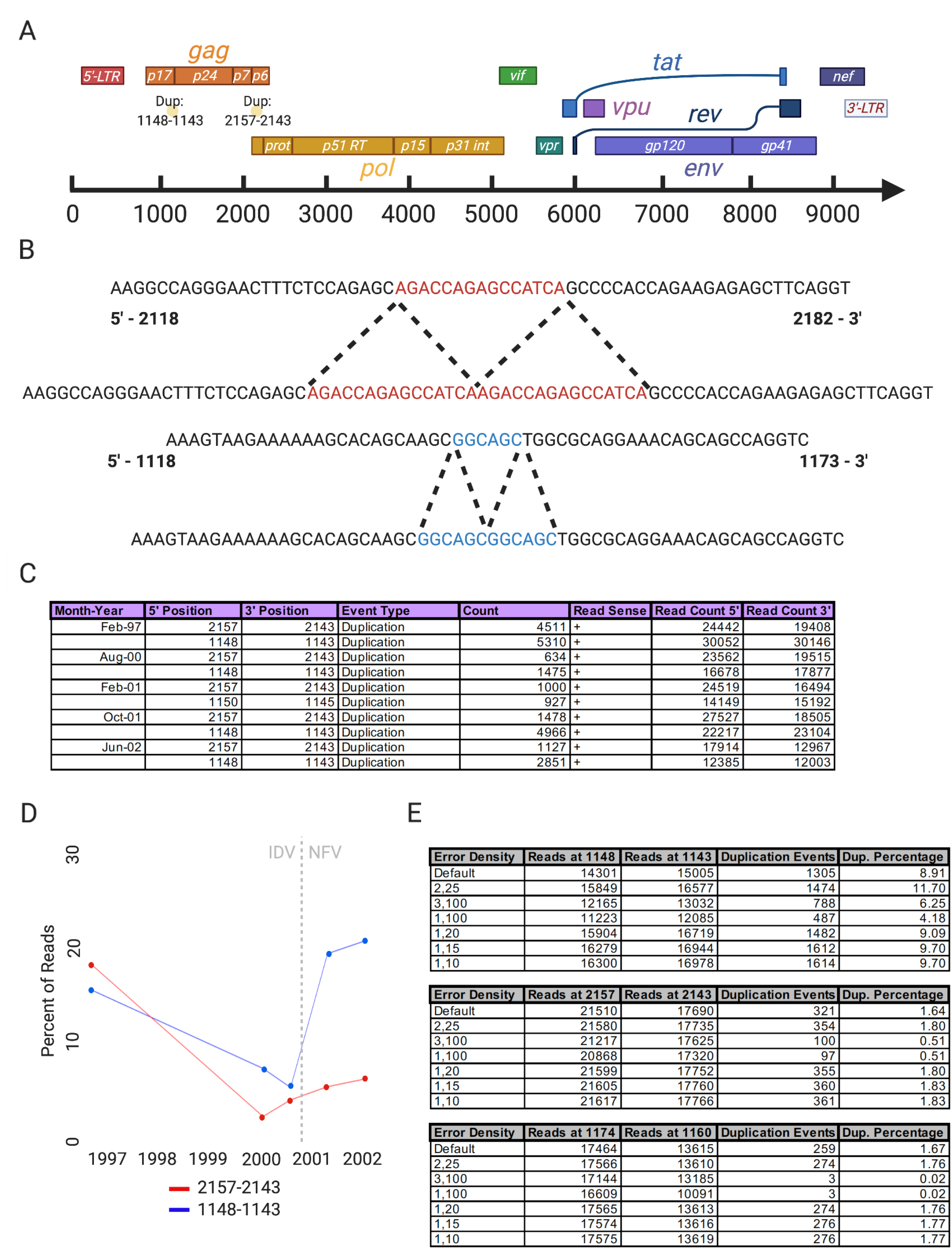
Duplications in *gag* in HIV patient data. **A)** Schematic of the HIV genome with the identified duplication events noted at nucleotide positions 1148-1143 and 2157-2143. **B)** Depiction of these duplication events at the sequence level. **C)** *ViReMa* output in the Virus_Recombination_Results.bed with all timepoints compiled and focused on the primary duplication events. **D)** Plot of the percent of mapped reads at these nucleotide positions that identify the noted duplication events over time. Note that the patient’s anti-retroviral treatment was changed after the third timepoint from indinavir (IDV) to nelfinavir (NFV). **E)** Tables depicting how the number of duplication events detected is dependent on the error density x,y where x is the number of allowed mismatches in y nucleotides. We include the two above mentioned events and an additional duplication event at position 1174.

We recently reported that single-nucleotide variants (SNVs) elsewhere in the viral genome are evolutionarily corelated with these duplications (57), some of which were found in close proximity to the duplication site (e.g. SNV D121A and Duplication event 1148^1143). Given the high intrahost diversity of the HIV population in these samples, careful consideration must be taken to the number of reference sequence mismatches that are tolerated during the *ViReMa* alignment process. If too few mismatches are allowed in each segment, then recombination events found near to SNVs will not be reported if the gap between the SNV and the recombination event is smaller than that selected mapping seed. We therefore used these data to investigate how changing the mismatch handling parameters affected the read alignment and recombination detection of *ViReMa*.

We mapped the Feb 2001 data using *ViReMa* with a range of error-handling parameters listed in **Figure 2E**. Using the ‘Error Density’ function allowing no more than two mismatches within a 25-nucleotide window yielded a small increase of ~10% in the number of mapped reads as well as a greater proportion of reported recombination events (~10% increase). The more permissive settings (1,10 and 1,15) increased the number of mapped reads by a further ~3%, but did not change the proportion of these that reported recombination events. The largest deviations in duplication events per reads at these particular nucleotide positions was found when using the least permissive parameters of 3,100 and 1,100, which as expected reduced both the number of reads mapped and the overall proportion of these reporting recombination events. This illustrates that a more permissive handling of mismatches surrounding putative recombination events can have an impact on how frequently they are reported. Altogether, these HIV patient datasets illustrate the ability of *ViReMa* to identify duplication events and how altering the permissibility of *ViReMa* by changing the error density parameter allows the detection of drug-resistant duplications that would otherwise have been missed due to their correlation to other drug-resistant SNVs.

### Copy-back DVGs are detected in the negative sense RNA virus, Sendai virus

Copy-back or snap-back RNAs are types of defective viral genomes (DVGs) commonly found in negative-sense RNA viruses including measles, mumps, Nipah and Sendai (SeV) virus (12). They emerge during synthesis of the viral genome (negative-sense) when using the anti-genome (positive-sense) as a template through a presumed mechanism in which the viral polymerase stalls and detaches from the template strand and begins copying the nascent strand creating a hairpin loop with complementary sequences in the strand at both ends. The existence of copy-bask DVGs is well characterized for SeV, (a paramyxovirus closely related to human parainfluenza viruses 1 and 3, also called murine parainfluenza virus 1) where sequencing found defective interfering genomes that consisted of the 5’ end of the genomic (negative sense) strand and the 3’ end of the positive sense or coding strand (61). Since their original description, these recombination events have been shown to be strongly immunostimulatory and to promote persistent viral infections by preventing apoptosis in DVG enriched cells versus highly infected cells with full-length virus (62).

As part of a previous study investigating the roles of copy-back DVGs in SeV pathogenesis (62), total RNA was purified from DVG-enriched infected cells in culture and used to synthesize cDNA libraries using an Illumina TruSeq Stranded Total RNA LT kit with Ribo-Zero Gold. cDNA libraries were sequenced on an Illumina NextSeq 550, obtaining 21-53M 75 bp single-end reads. Here, we focused on the SeV sample experimentally determined to contain a high abundance of one species of copy-back DVG. We mapped these reads using *ViReMa* to both the SeV genome (AB855654.1) and the human genome (hg19) to confirm the presence of known copy-back DVGs (**Figure 3**). *ViReMa* detected 21 unique potential copy-back DVGs in this dataset (**SData 3**), however only four of these were detected with more than one mapped read. The two most common events were negative to positive sense fusions of the SeV genome at coordinates 15291^(-)^^14933^(+)^ and 14932^(-)^^15292^(+)^ as illustrated in **Figure 3B,C,** with 50,574 and 3,611 reads respectively. This major copy-back DVG was also identified using VODKA (63). These read counts are in far excess of the number of reads found for other types of RNA recombination events such as deletion-type event, with the most abundance recombination event having only 29 reads (**SData 3**).

**Figure 3.**
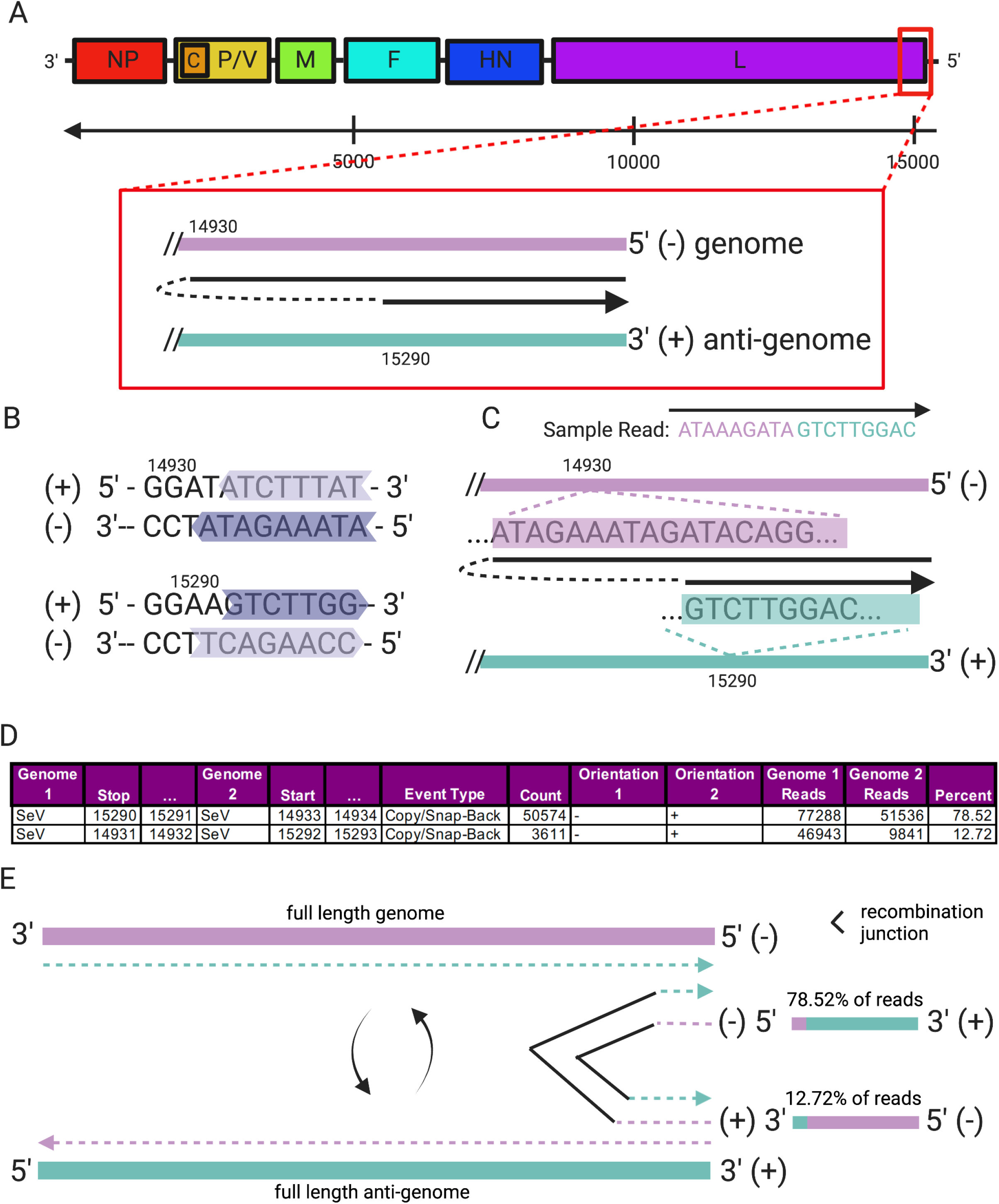
Detection of copy-back RNAs in Sendai virus (SeV) using *ViReMa*. **A)** Map of the negative sense (packaged) SeV genome with a box around the 5’ end where copy-back DVGs occur. Expanded in the box is a model for how copy-backs are made by template switching of the viral polymerase. **B)** Sequences in the positive and negative sense strands of SeV where template switching produces copy-back RNAs. Here the arrow of each color indicates direction of the polymerase and the color indicates a particular copy-back, ex. light purple and purple are separate copy-backs. **C)** Depiction of a sample read identifying a copy-back and how that aligns to the 5’ end of the negative sense strand and 3’ end of the positive sense strand. **D)** Table showing the number of copy-back events in the data set. The ‘Percent’ column is calculated from the count of reads with the recombination event (from Virus_Fusions.BEDPE) compared to the total number of reads at those nucleotide positions (calculated using *bedtools*). **E)** Our results indicate the presence of a secondary copy-back genome. While the primary copy-back is produced by a template switching event from a negative (genome) sense to positive (anti-genome) sense template, the secondary copy-back is produced by replication using the primary copy-back as a template. The secondary copy-back accounts for 78% of reads at these genome positions (15290 [-] and 14933 [+]).

The data in the table in **Figure 3D** is derived from the *ViReMa* output file ‘*Virus_Fusions.BEDPE*’. The ‘Reads’ columns describe the number of reads at each particular nucleotide position of SeV and the ‘Percent’ column was populated by calculating the number of recombination event counts over the average of the 3’ and 5’ read counts. Although reported separately, these are the same copy-back DVG but reflect reads mapping to both the sense and the anti-sense version of this DVG. Interestingly, although 14932^(-)^^15292^(+)^ is the junction that would be expected in the original copy-back DVG generated during synthesis from the positive-sense anti-genome, the reverse-complementary 15291^(-)^^14933^(+)^ events is approximately 6.5-fold more abundant. As the cDNA synthesis strategy used retains the original stranded-ness of the input RNA, this data therefore suggests that multiple anti-sense DVGs are generated from the original DVG. This example demonstrates the ability of *ViReMa* to faithfully identify copy-back recombination events and use their relative abundance to reveal that DVGs are actively replicated into anti-DVGs.

### Sulfolobus turreted icosahedral virus (STIV) captures fragments of the host genome

*Sulfolobus* turreted icosahedral virus (STIV) infects the thermophile *Sulfolobus solfataricus* (SSP2). These archaea, and the viruses that infect them, are found in hot springs and have evolved to withstand their harsh environments. Virus particles are 74 nm in diameter, contain an inner membrane within its icosahedral capsid, and enclose a 17.7 kb double-stranded circular DNA genome (64, 65). To characterize rates of recombination in a DNA virus, we obtained genomic DNA from purified particles of STIV expressed in culture (66). Genomic viral DNA was prepared for NGS using ClickSeq (39) and sequenced on an Illumina HiSeq 1000 obtaining ~36M single-end 150 bp reads. We used *ViReMa* to map these reads to the STIV genome (NC_005892.1) and the host genome (*Sulfolobus*: AE006641.1) in parallel (BED files are provided in **SData 4**). As expected, the majority of the NGS reads mapped directly to the virus genome (**Table 1**). Interestingly however, we also observed a large number of ‘virus-to-host’ recombination events. Such recombination events are illustrated in **Figure 4A** and the output information reported in the BEDPE file is shown in **Figure 4B**. The column labeled ‘Reads’ describes the number of reads at that nucleotide position of STIV (obtained using *bedtools*) and the final column was added to highlight the percentage of viral reads at that position containing that particular recombination event.

**Figure 4.**
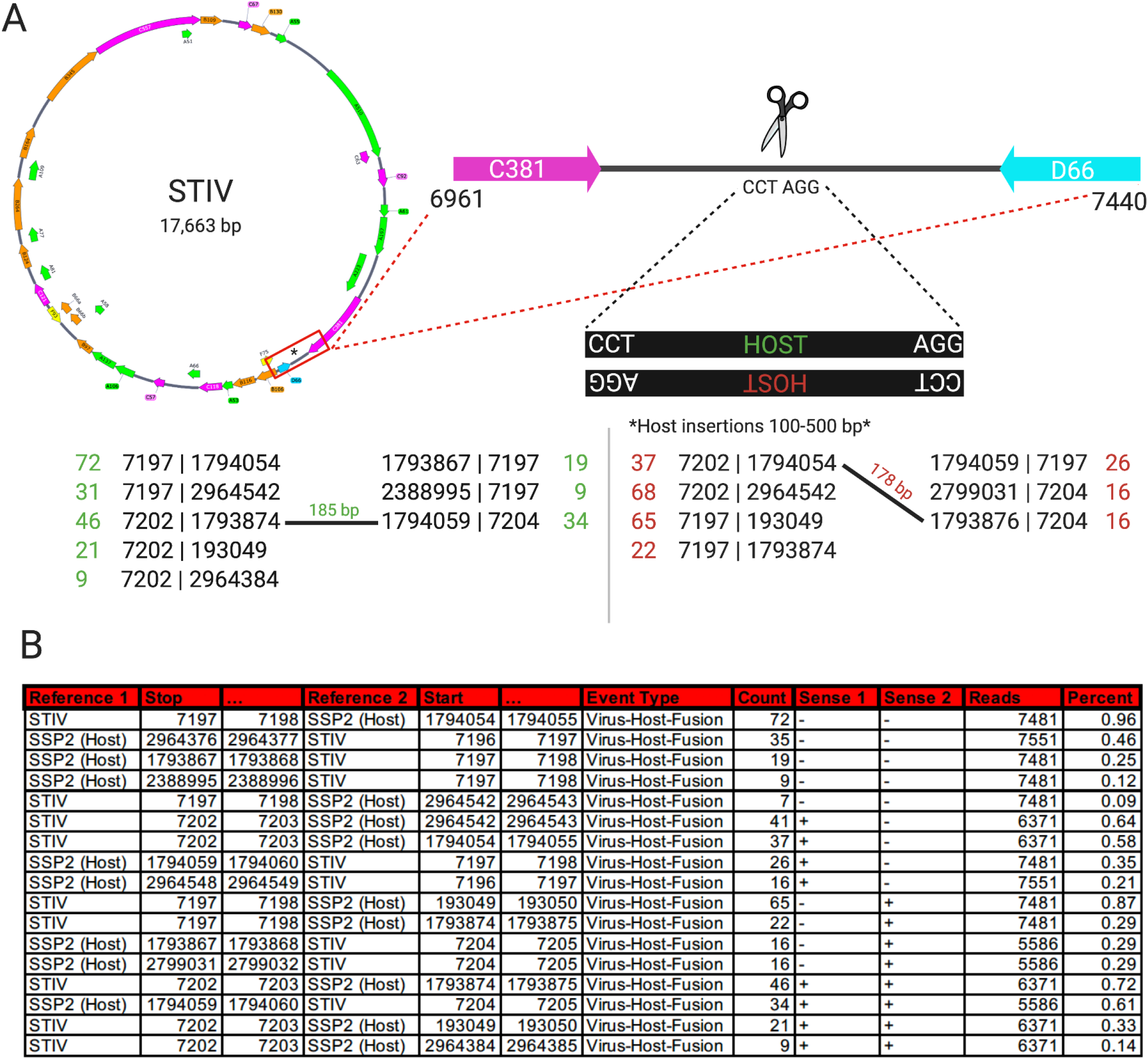
Host-to-virus recombination in *Sulfolobus* turreted icosahedral virus (STIV) as detected by *ViReMa*. **A)** Map of the circular STIV genome and a zoomed in region between genes C381 and D66 with a CCTAGG motif where host-to-virus recombination occurs. We list the recombination events found in the dataset at that site. Here, upside-down ‘HOST’ in red is from the opposite strand of STIV, and therefore recombination occurs at the same site and the motif is in reverse complement. **B)** ViReMa output from the Virus-to-Host_Recombinations.BEDPE. Where ‘Reference 1’ is the first genome mapped to a read up until position ‘Stop’, ‘Reference 2’ is the second genome mapped from ‘Start’ and ‘Sense 1’/’Sense 2’ refers to the sense of the respective reference genome. The ‘Count’ is the number of reads containing the fusion event and ‘Reads’ is the total number of reads mapped to the virus at that position (calculated using *bedtools*). ‘Percent’ was calculated by comparing the ‘Count’ to ‘Reads’ at that position.

**Table 1:** STIV mapping Statistics.

|  |  |
| --- | --- |
| Total Reads | 36,298,490 |
| STIV | 35,831,428 |
| SSP2 | 8,598 |
| Unmapped | 458,464 |
| Virus Recombination Events | 38,150 |
| Host Recombination Events | 35 |
| Virus-to-Host Recombination | 1,770 |

The recombination junction breakpoints in the viral genome were strongly enriched at nucleotide position 7200 of the STIV genome, which is in a noncoding region between the genes encoding viral proteins C381 and D66 (**Figure 4A**). The recombination junction breakpoints were found at a few different loci within the host genome. Reads mapping over recombination junctions are found in both positive and negative sense orientations. Furthermore, many of the host recombination junctions are occurred very close to one another (pairs of junctions are observed within 100-500nts of each other. These observations are all consistent with the insertion of small (100-500 nt) portions of host genome into the viral genome in both positive and negative orientations (illustrated in **Figure 4A**). Interestingly, both the viral loci for these recombination junctions, and all the junction sites of the inserted host segments, occur at ‘CCTAGG’ motifs. The palindromic nature of this motif is reminiscent of mechanisms employed by bacteriophages that utilize recombinases to cut and transpose the phage genome into the host genome at short complementary palindromic sites to form a lysogenic prophage. STIV has previously been shown to undergo homologous recombination with various species of *Sulfolobus* at sites with this motifs (67).

We posit a mechanism akin to classical bacterial transduction, whereby STIV integrates its own genome into the host genome at ‘CCTAGG’ sites using a viral recombinase. Upon reactivation, the CCTAGG sites at either end of the viral prophage are re-cut to release the viral prophage. However, if there is an additional CCTAGG site in the host genome that is close to the prophage integration site, then a small additional segment of host DNA is inadvertently introduced into the viral genome as a small insertion. As the Illumina reads are too-short (150nts) to completely map across these small host DNA insertions and into the flanking viral genome on both sides of the insertion, it is also possible that these recombination events correspond to the virus-to-host junctions of proviral integration sites into the host genome. However, the input DNA was obtained from purified virus particles expressed in culture. These data therefore demonstrate the ability of ViReMa to identify virus-to-host recombination events and reveals a putative capture of host DNA into a viral vector as a consequence of proviral integration.

## Discussion

*ViReMa* has provided a versatile tool for the detection and enumeration of recombination events in a broad array of virus families. *ViReMa* has been independently validated (33, 34) demonstrating robust sensitivity and accuracy. To keep up with continuous improvements in Next-Generation Sequencing platforms (such as longer read lengths) and to accurately capture recombination events found within complex and diverse populations of virus genomes, we present here a renovated handling of mismatches in aligned read segments. We also provide standardized outputs (SAM, BED and BEDPE files) for both the read alignment and discovered recombination junctions that allow integration of the outputs of *ViReMa* with other sequence visualization software and bioinformatic packages. Using a series of ‘case studies’ we demonstrate how *ViReMa* can be employed to capture different recombination events. These include deletions in FHV; short duplications in the HIV genome, copy-back RNAs in SeV; as well as virus-to-host fusion events in STIV.

*ViReMa* strictly requires a user-defined number of nucleotides to be mapped either side of a putative recombination junction, providing confidence in the identity of the recombination event reported, though at the expense of sensitivity. For situations where the predominant form of recombination event is a deletion, canonical pipelines for splice detection such as HISAT2 (21) and STAR (23) would provide a quicker and more sensitive route to detect these events as they do not rely on multiple iterations of *bowtie* alignment and use a dynamic seed-based heuristic. Additionally, where recombination/deletion events are known *a priori*, annotated junction events can be utilized in the index building steps of HISAT2 (21) and *STAR* (23). In these cases however, recombination acceptor sites must be located downstream of recombination donor sites within the same single reference sequence, as would be the case in canonical eukaryotic splicing events. Additionally, these aligners may be overly permissive in requiring mapped nucleotides on either side of a recombination junction resulting in the reporting of artifactual events. For eukaryotic splicing, this permissibility is acceptable as splice events are restricted to a small number of possible sites. However, this is not the case for viral recombination where junctions are often diverse and unpredictable, as previously noted for RNA viruses such as FHV (68). To reflect this, *ViReMa* enforces strict parameters for read mapping either side of a junction for all identified recombination events.

Recently, due to the increased interest in copy-back and snap-back defective viral genomes (DVGs) in negative-strand RNA viruses such as Respiratory Syncytial virus (69), a number of platforms have been proposed to specifically map these species including DI-Tector (26) and VODKA (27). Early versions of DI-Tector (26) and VODKA (27) search for negative-strand to positive-strand fusion events which are found in the copy-back and snap-back DVGs. The VODKA algorithm will generate numerous reference sequences based upon putative copy/snap-back viral genome rearrangements and align short reads to a ‘pseudo-library’. This requires prior information to focus the pseudo-reference library to regions suspected or known to form DVGs. Without this knowledge, the pseudo-reference library can become exponentially large with increased genome size. However, by providing a range of putative sequences with pre-defined recombination junctions, this constitutes a more sensitive approach as unambiguous mappings can be found across these junctions’ sites whilst requiring a smaller seed region. Further, search-seed lengths and tolerance to non-reference variations can be optimized for this application. Therefore, similar to finding known deletion/splice-like events using *HISAT2/STAR* aligners, in scenarios where information about the types of putative the copy/snap-back DVGs is known, VODKA provides a more sensitive readout. In contrast, when no prior information is available, *ViReMa* provides an ‘agnostic’ approach, but at the expense of sensitivity.

Currently, ViReMa has been validated and tested solely on high-accuracy short-read Next-Generation Sequencing data. With continued improvements to long-read sequencing platforms in terms of data yield, sequence length and accuracy (such as the Oxford Nanopore Technologies R10.4 nanopore), future applications of *ViReMa* may include the discovery of recombination or other types of fusion events in nucleic acids not restricted to viral species. Overall, the broad utility of *ViReMa* derives from its strategy whereby read segments are mapped entirely independently from other read segments. This versatility places it as useful all-in-one tool, particularly when *a priori* knowledge is lacking, for the discovery, mapping and annotations of viral recombination events.

## Supporting information

SData1

SData2

SData3

SData4

## Acknowledgements

We would like to thank Dr. Carolina Lopez (Washington University School of Medicine in St. Louis) for discussions and critical reading of the manuscript and providing the SeV RNA. We would like to thank Dr. Mark Young (Montana State University) for providing viral DNA from purified STIV particles. This work was funded by: NIH grant R21AI151725 from to ALR; National Institute of Allergy and Infectious Diseases U54AI150472 to BET and ALR; CDC contract 200-2021-11195 to ALR; sub-contract from NIH grant R01AI042189 to ALR. SLS is supported by a Kempner fellowship.

---

**Supplementary Data 1:** Output files in BED format from ViReMa analysis of simulated Flock house virus (FHV) data.

**Supplementary Data 2:** Output files in BED format from ViReMa analysis of clinical specimens of HIV across five time-points.

**Supplementary Data 3:** Output files in BED format from ViReMa analysis of purified Sendai virus (SeV).

**Supplementary Data 4:** Output files in BED format from ViReMa analysis of Sulfolobus Turreted Icosahedral Virus (STIV).

## References

1. Simon-Loriere E, Holmes EC. 2011. Why do RNA viruses recombine? Nat Rev Microbiol 9:617–26.

2. Li X, Giorgi EE, Marichannegowda MH, Foley B, Xiao C, Kong XP, Chen Y, Gnanakaran S, Korber B, Gao F. 2020. Emergence of SARS-CoV-2 through recombination and strong purifying selection. Sci Adv 6.

3. Palmenberg AC, Spiro D, Kuzmickas R, Wang S, Djikeng A, Rathe JA, Fraser-Liggett CM, Liggett SB. 2009. Sequencing and analyses of all known human rhinovirus genomes reveal structure and evolution. Science 324:55–9.

4. Lau SK, Feng Y, Chen H, Luk HK, Yang WH, Li KS, Zhang YZ, Huang Y, Song ZZ, Chow WN, Fan RY, Ahmed SS, Yeung HC, Lam CS, Cai JP, Wong SS, Chan JF, Yuen KY, Zhang HL, Woo PC. 2015. Severe Acute Respiratory Syndrome (SARS) Coronavirus ORF8 Protein Is Acquired from SARS-Related Coronavirus from Greater Horseshoe Bats through Recombination. J Virol 89:10532–47.

5. Johnson BA, Xie X, Bailey AL, Kalveram B, Lokugamage KG, Muruato A, Zou J, Zhang X, Juelich T, Smith JK, Zhang L, Bopp N, Schindewolf C, Vu M, Vanderheiden A, Winkler ES, Swetnam D, Plante JA, Aguilar P, Plante KS, Popov V, Lee B, Weaver SC, Suthar MS, Routh AL, Ren P, Ku Z, An Z, Debbink K, Diamond MS, Shi PY, Freiberg AN, Menachery VD. 2021. Loss of furin cleavage site attenuates SARS-CoV-2 pathogenesis. Nature 591:293–299.

6. Muth D, Corman VM, Roth H, Binger T, Dijkman R, Gottula LT, Gloza-Rausch F, Balboni A, Battilani M, Rihtaric D, Toplak I, Ameneiros RS, Pfeifer A, Thiel V, Drexler JF, Muller MA, Drosten C. 2018. Attenuation of replication by a 29 nucleotide deletion in SARS-coronavirus acquired during the early stages of human-to-human transmission. Sci Rep 8:15177.

7. Wang S, Sotcheff SL, Gallardo CM, Jaworski E, Torbett BE, Routh AL. 2021. Co-variation of viral recombination with single nucleotide variants during virus evolution revealed by CoVaMa. bioRxiv doi:10.1101/2021.09.14.460373:2021.09.14.460373.

8. Martins AN, Waheed AA, Ablan SD, Huang W, Newton A, Petropoulos CJ, Brindeiro RD, Freed EO. 2016. Elucidation of the Molecular Mechanism Driving Duplication of the HIV-1 PTAP Late Domain. J Virol 90:768–79.

9. Gallardo CM, Wang S, Montiel-Garcia DJ, Little SJ, Smith DM, Routh AL, Torbett BE. 2021. MrHAMER yields highly accurate single molecule viral sequences enabling analysis of intra-host evolution. Nucleic Acids Res doi:10.1093/nar/gkab231.

10. Tamiya S, Mardy S, Kavlick MF, Yoshimura K, Mistuya H. 2004. Amino acid insertions near Gag cleavage sites restore the otherwise compromised replication of human immunodeficiency virus type 1 variants resistant to protease inhibitors. J Virol 78:12030–40.

11. Peters S, Munoz M, Yerly S, Sanchez-Merino V, Lopez-Galindez C, Perrin L, Larder B, Cmarko D, Fakan S, Meylan P, Telenti A. 2001. Resistance to nucleoside analog reverse transcriptase inhibitors mediated by human immunodeficiency virus type 1 p6 protein. J Virol 75:9644–53.

12. Genoyer E, Lopez CB. 2019. The Impact of Defective Viruses on Infection and Immunity. Annu Rev Virol doi:10.1146/annurev-virology-092818-015652.

13. Vignuzzi M, Lopez CB. 2019. Defective viral genomes are key drivers of the virus-host interaction. Nat Microbiol doi:10.1038/s41564-019-0465-y.

14. Huang AS, Baltimore D. 1970. Defective viral particles and viral disease processes. Nature 226:325–7.

15. Dimmock NJ, Easton AJ. 2014. Defective interfering influenza virus RNAs: time to reevaluate their clinical potential as broad-spectrum antivirals? J Virol 88:5217–27.

16. Alnaji FG, Brooke CB. 2020. Influenza virus DI particles: Defective interfering or delightfully interesting? PLoS Pathog 16:e1008436.

17. Felt SA, Sun Y, Jozwik A, Paras A, Habibi MS, Nickle D, Anderson L, Achouri E, Feemster KA, Cardenas AM, Turi KN, Chang M, Hartert TV, Sengupta S, Chiu C, Lopez CB. 2021. Detection of respiratory syncytial virus defective genomes in nasal secretions is associated with distinct clinical outcomes. Nat Microbiol 6:672–681.

18. Albarino CG, Price BD, Eckerle LD, Ball LA. 2001. Characterization and template properties of RNA dimers generated during flock house virus RNA replication. Virology 289:269–82.

19. Nguyen HT, Torian U, Faulk K, Mather K, Engle RE, Thompson E, Bonkovsky HL, Emerson SU. 2012. A naturally occurring human/hepatitis E recombinant virus predominates in serum but not in faeces of a chronic hepatitis E patient and has a growth advantage in cell culture. J Gen Virol 93:526–530.

20. Shukla P, Nguyen HT, Faulk K, Mather K, Torian U, Engle RE, Emerson SU. 2012. Adaptation of a genotype 3 hepatitis E virus to efficient growth in cell culture depends on an inserted human gene segment acquired by recombination. J Virol 86:5697–707.

21. Kim D, Langmead B, Salzberg SL. 2015. HISAT: a fast spliced aligner with low memory requirements. Nat Methods 12:357–60.

22. Pertea M, Kim D, Pertea GM, Leek JT, Salzberg SL. 2016. Transcript-level expression analysis of RNA-seq experiments with HISAT, StringTie and Ballgown. Nat Protoc 11:1650–67.

23. Dobin A, Davis CA, Schlesinger F, Drenkow J, Zaleski C, Jha S, Batut P, Chaisson M, Gingeras TR. 2013. STAR: ultrafast universal RNA-seq aligner. Bioinformatics 29:15–21.

24. Li JW, Wan R, Yu CS, Co NN, Wong N, Chan TF. 2013. ViralFusionSeq: accurately discover viral integration events and reconstruct fusion transcripts at single-base resolution. Bioinformatics 29:649–51.

25. Wang Q, Jia P, Zhao Z. 2015. VERSE: a novel approach to detect virus integration in host genomes through reference genome customization. Genome Med 7:2.

26. Beauclair G, Mura M, Combredet C, Tangy F, Jouvenet N, Komarova AV. 2018. DI-tector: defective interfering viral genomes’ detector for next-generation sequencing data. Rna 24:1285–1296.

27. Sun Y, Kim E, Speranza E, Taylor L, Agarwal D, Gerhardt D, Bennett R, Connor J, Grant G, Lopez C. 2018. doi:10.1101/349001.

28. Karagiannis K, Simonyan V, Chumakov K, Mazumder R. 2017. Separation and assembly of deep sequencing data into discrete sub-population genomes. Nucleic Acids Res 45:10989–11003.

29. Wood GR, Ryabov EV, Fannon JM, Moore JD, Evans DJ, Burroughs N. 2014. MosaicSolver: a tool for determining recombinants of viral genomes from pileup data. Nucleic Acids Res 42:e123.

30. Routh A, Johnson JE. 2014. Discovery of functional genomic motifs in viruses with ViReMa-a Virus Recombination Mapper-for analysis of next-generation sequencing data. Nucleic Acids Res 42:e11.

31. Langmead B, Trapnell C, Pop M, Salzberg SL. 2009. Ultrafast and memory-efficient alignment of short DNA sequences to the human genome. Genome Biol 10:R25.

32. Li H, Durbin R. 2009. Fast and accurate short read alignment with Burrows-Wheeler transform. Bioinformatics 25:1754–60.

33. Alnaji FG, Holmes JR, Rendon G, Vera JC, Fields CJ, Martin BE, Brooke CB. 2019. Sequencing Framework for the Sensitive Detection and Precise Mapping of Defective Interfering Particle-Associated Deletions across Influenza A and B Viruses. J Virol 93.

34. Boussier J, Munier S, Achouri E, Meyer B, Crescenzo-Chaigne B, Behillil S, Enouf V, Vignuzzi M, van der Werf S, Naffakh N. 2020. RNA-seq accuracy and reproducibility for the mapping and quantification of influenza defective viral genomes. RNA 26:1905–1918.

35. Bertran A, Ciuffo M, Margaria P, Rosa C, Oliveira Resende R, Turina M. 2016. Host-specific accumulation and temperature effects on the generation of dimeric viral RNA species derived from the S-RNA of members of the Tospovirus genus. J Gen Virol 97:3051–3062.

36. Xu C, Sun X, Taylor A, Jiao C, Xu Y, Cai X, Wang X, Ge C, Pan G, Wang Q, Fei Z, Wang Q. 2017. Diversity, Distribution, and Evolution of Tomato Viruses in China Uncovered by Small RNA Sequencing. J Virol 91.

37. Kutnjak D, Rupar M, Gutierrez-Aguirre I, Curk T, Kreuze JF, Ravnikar M. 2015. Deep sequencing of virus-derived small interfering RNAs and RNA from viral particles shows highly similar mutational landscapes of a plant virus population. J Virol 89:4760–9.

38. Jaworski E, Routh A. 2017. Parallel ClickSeq and Nanopore sequencing elucidates the rapid evolution of defective-interfering RNAs in Flock House virus. PLoS Pathog 13:e1006365.

39. Routh A, Head SR, Ordoukhanian P, Johnson JE. 2015. ClickSeq: Fragmentation-Free Next-Generation Sequencing via Click Ligation of Adaptors to Stochastically Terminated 3’-Azido cDNAs. J Mol Biol 427:2610–6.

40. Barrows NJ, Campos RK, Powell ST, Prasanth KR, Schott-Lerner G, Soto-Acosta R, Galarza-Muñoz G, McGrath EL, Urrabaz-Garza R, Gao J, Wu P, Menon R, Saade G, Fernandez-Salas I, Rossi SL, Vasilakis N, Routh A, Bradrick SS, Garcia-Blanco MA. 2016. A Screen of FDA-Approved Drugs for Inhibitors of Zika Virus Infection. Cell Host Microbe 20:259–70.

41. Langsjoen RM, Zhou Y, Holcomb RJ, Routh AL. 2021. Chikungunya Virus Infects the Heart and Induces Heart-Specific Transcriptional Changes in an Immunodeficient Mouse Model of Infection. Am J Trop Med Hyg doi:10.4269/ajtmh.21-0719.

42. Langsjoen RM, Muruato AE, Kunkel SR, Jaworski E, Routh A. 2020. Differential Alphavirus Defective RNA Diversity between Intracellular and Extracellular Compartments Is Driven by Subgenomic Recombination Events. mBio 11.

43. Alnaji FG, Holmes JR, Rendon G, Vera JC, Fields CJ, Martin BE, Brooke CB. 2018. doi:10.1101/440651.

44. Smith SC, Gribble J, Diller JR, Wiebe MA, Thoner TW, Jr., Denison MR, Ogden KM. 2021. Reovirus RNA recombination is sequence directed and generates internally deleted defective genome segments during passage. J Virol doi:10.1128/JVI.02181-20.

45. Gribble J, Stevens LJ, Agostini ML, Anderson-Daniels J, Chappell JD, Lu X, Pruijssers AJ, Routh AL, Denison MR. 2021. The coronavirus proofreading exoribonuclease mediates extensive viral recombination. PLoS Pathog 17:e1009226.

46. Jaworski E, Langsjoen RM, Mitchell B, Judy B, Newman P, Plante JA, Plante KS, Miller AL, Zhou Y, Swetnam D, Sotcheff S, Morris V, Saada N, Machado RR, McConnell A, Widen SG, Thompson J, Dong J, Ren P, Pyles RB, Ksiazek TG, Menachery VD, Weaver SC, Routh AL. 2021. Tiled-ClickSeq for targeted sequencing of complete coronavirus genomes with simultaneous capture of RNA recombination and minority variants. Elife 10.

47. Li H, Handsaker B, Wysoker A, Fennell T, Ruan J, Homer N, Marth G, Abecasis G, Durbin R. 2009. The Sequence Alignment/Map format and SAMtools. Bioinformatics 25:2078–9.

48. Quinlan AR, Hall IM. 2010. BEDTools: a flexible suite of utilities for comparing genomic features. Bioinformatics 26:841–2.

49. Langmead B, Salzberg SL. 2012. Fast gapped-read alignment with Bowtie 2. Nat Methods 9:357–9.

50. Jo H, Koh G. 2015. Faster single-end alignment generation utilizing multi-thread for BWA. Biomed Mater Eng 26 Suppl 1:S1791–6.

51. Milne I, Bayer M, Cardle L, Shaw P, Stephen G, Wright F, Marshall D. 2010. Tablet--next generation sequence assembly visualization. Bioinformatics 26:401–2.

52. Thorvaldsdottir H, Robinson JT, Mesirov JP. 2013. Integrative Genomics Viewer (IGV): high-performance genomics data visualization and exploration. Brief Bioinform 14:178–92.

53. Robinson JT, Thorvaldsdottir H, Winckler W, Guttman M, Lander ES, Getz G, Mesirov JP. 2011. Integrative genomics viewer. Nat Biotechnol 29:24–6.

54. Huang W, Li L, Myers JR, Marth GT. 2012. ART: a next-generation sequencing read simulator. Bioinformatics 28:593–4.

55. Genomes Project C, Auton A, Brooks LD, Durbin RM, Garrison EP, Kang HM, Korbel JO, Marchini JL, McCarthy S, McVean GA, Abecasis GR. 2015. A global reference for human genetic variation. Nature 526:68–74.

56. Routh A, Chang MW, Okulicz JF, Johnson JE, Torbett BE. 2015. CoVaMa: Co-Variation Mapper for disequilibrium analysis of mutant loci in viral populations using next-generation sequence data. Methods doi:10.1016/j.ymeth.2015.09.021.

57. Wang S, Sotcheff SL, Gallardo CM, Jaworski E, Torbett BE, Routh AL. 2022. Covariation of viral recombination with single nucleotide variants during virus evolution revealed by CoVaMa. Nucleic Acids Res doi:10.1093/nar/gkab1259.

58. Flynn WF, Chang MW, Tan Z, Oliveira G, Yuan J, Okulicz JF, Torbett BE, Levy RM. 2015. Deep sequencing of protease inhibitor resistant HIV patient isolates reveals patterns of correlated mutations in Gag and protease. PLoS Comput Biol 11:e1004249.

59. Wu TD, Schiffer CA, Gonzales MJ, Taylor J, Kantor R, Chou S, Israelski D, Zolopa AR, Fessel WJ, Shafer RW. 2003. Mutation patterns and structural correlates in human immunodeficiency virus type 1 protease following different protease inhibitor treatments. J Virol 77:4836–47.

60. Rhee SY, Liu TF, Holmes SP, Shafer RW. 2007. HIV-1 subtype B protease and reverse transcriptase amino acid covariation. PLoS Comput Biol 3:e87.

61. Re GG, Gupta KC, Kingsbury DW. 1983. Sequence of the 5’ end of the Sendai virus genome and its variable representation in complementary form at the 3’ ends of copy-back defective interfering RNA species: identification of the L gene terminus. Virology 130:390–6.

62. Xu J, Sun Y, Li Y, Ruthel G, Weiss SR, Raj A, Beiting D, Lopez CB. 2017. Replication defective viral genomes exploit a cellular pro-survival mechanism to establish paramyxovirus persistence. Nat Commun 8:799.

63. Sun Y, Kim EJ, Felt SA, Taylor LJ, Agarwal D, Grant GR, Lopez CB. 2019. A specific sequence in the genome of respiratory syncytial virus regulates the generation of copy-back defective viral genomes. PLoS Pathog 15:e1007707.

64. Rice G, Tang L, Stedman K, Roberto F, Spuhler J, Gillitzer E, Johnson JE, Douglas T, Young M. 2004. The structure of a thermophilic archaeal virus shows a double-stranded DNA viral capsid type that spans all domains of life. Proc Natl Acad Sci U S A 101:7716–20.

65. Veesler D, Ng TS, Sendamarai AK, Eilers BJ, Lawrence CM, Lok SM, Young MJ, Johnson JE, Fu CY. 2013. Atomic structure of the 75 MDa extremophile Sulfolobus turreted icosahedral virus determined by CryoEM and X-ray crystallography. Proc Natl Acad Sci U S A 110:5504–9.

66. Fu CY, Wang K, Gan L, Lanman J, Khayat R, Young MJ, Jensen GJ, Doerschuk PC, Johnson JE. 2010. In vivo assembly of an archaeal virus studied with whole-cell electron cryotomography. Structure 18:1579–86.

67. Mao D, Grogan DW. 2017. How a Genetically Stable Extremophile Evolves: Modes of Genome Diversification in the Archaeon Sulfolobus acidocaldarius. J Bacteriol 199.

68. Jaworski E, Routh A. 2017. Parallel ClickSeq and Nanopore Sequencing elucidates the rapid evolution of Defective-Interfering RNAs in Flock House virus. PLOS Pathogens.

69. Sun Y, Jain D, Koziol-White CJ, Genoyer E, Gilbert M, Tapia K, Panettieri RA, Hodinka RL, López CB. 2015. Immunostimulatory Defective Viral Genomes from Respiratory Syncytial Virus Promote a Strong Innate Antiviral Response during Infection in Mice and Humans. PLoS Pathog 11:e1005122.

